# Endogenous retrovirus ERV-DC8 highly integrated in domestic cat populations is a replication-competent provirus

**DOI:** 10.1101/2024.05.09.593269

**Authors:** Didik Pramono, Yutaro Muto, Yo Shimazu, R.M.C. Deshapriya, Isaac Makundi, MaríaCruz Arnal, Daniel Fernández de Luco, Minh Ha Ngo, Ariko Miyake, Kazuo Nishigaki

## Abstract

Endogenous retroviruses (ERVs) are remnants of ancient retroviral infections in vertebrate genomes and are inherited by offspring. ERVs can produce pathogenic viruses through gene mutations or recombination. ERVs in domestic cats (ERV-DCs) generate feline leukemia virus subgroup D (FeLV-D) through viral recombination. Herein, we characterized the locus ERV-DC8, on chromosome B1, as an infectious replication-competent provirus. ERV-DC8 infected several cell lines, including human cells. Transmission electron microscopy of ERV-DC8 identified the viral release as a *Gammaretrovirus*. ERV-DC8 was identified as the FeLV-D viral interference group, with feline copper transporter 1 as its viral receptor. Insertional polymorphism analysis showed high ERV-DC8 integration in domestic cats. This study highlights the role, pathogenicity, and evolutionary relationships between ERVs and their hosts.

## Introduction

Endogenous retroviruses (ERVs) are remnants of ancient retroviral infections and comprise approximately 4-10% of the human, mouse, and cat genomes (1-3). The invasion of germ cells causes endogenization via vertical transmission of the provirus to the offspring. Numerous ERVs are inactivated by the accumulation of mutations such as nucleotide sequence substitutions, insertions, and deletions, and/or by the loss of transcriptional activity due to epigenetic control (4, 5). These events may be related to viral integration time. Therefore, most ERVs exist as non-functional DNA. However, some endogenous retroviruses retain their infectivity and ability to replicate (6, 7). Additionally, both pathogenic and emerging viruses are generated by gene mutations or recombination (8). Various diseases associated with ERV have been reported in mice and humans (9, 10).

The feline leukemia virus (FeLV) is a pathogenic *Gammaretrovirus* that causes various pathological conditions in domestic cats, including proliferative and immune diseases. FeLV subgroups are classified according to viral entry based on receptor differences. FeLV subgroup A (FeLV-A) is the most common and is horizontally transmitted between hosts via grooming or biting (11). A novel FeLV, namely FeLV subgroup D (FeLV-D), has been identified in cases of feline lymphoma and leukemia (7, 12). FeLV-D was generated by the recombination of ERV of domestic cats (ERV-DCs) genotype I (GI) in the *env* region with FeLV (7), demonstrated a distinct tropism compared with other identified FeLV subgroups (7) and belonged to a different interference group. In addition, recombination between ERV-DCs and baboon endogenous retrovirus (BaEV) generated RD-114, another distinct feline retrovirus. This recombination event supports interspecies retroviral transmission between cats and primates (7, 13-15) and reinforces that the interactions between exogenous retroviruses and ERVs contribute to long-term retroviral diversification and evolution.

ERV-DC is an endogenous *Gammaretrovirus* of the domestic cat. In the last decade, ERV-DCs have been characterized and identified as young ERVs, estimated to have integrated into the host genome approximately 2.8 million years ago (MYA) (7). Thirteen loci of ERV-DC have been reported in the cat (*Felis silvestris catus*) genome (7). ERV-DCs are phylogenetically classified into genotypes I–III (GI, GII, and GIII). ERV-DCs typically have simple retroviral structures consisting of *gag, pol*, and *env* genes flanked by two long terminal repeats (LTRs) approximately 9 kb in length in the viral genome (7). Among the 13 ERV-DCs, ERV-DC8 (GI), ERV-DC10 (GIII), ERV-DC14 (GI), and ERV-DC18 (GIII) have intact open reading frames (ORFs). ERV-DC10, ERV-DC14, and ERV-DC18 can produce infectious viral particles that are characteristic of pantropic and xenotropic viruses (7). However, whether ERV-DC8 is replication-competent remains unclear. Notably, ERV-DCs are known to retain their ability to potentially affect their hosts via viral activity and contribute to the emergence of recombinant viruses. Here, we characterized, from a cat sample, ERV-DC8, its ability to infect various host cells, and its integration frequency in domestic cats.

## Materials and Methods

### Animals and Sampling

We used the DNA genome of the domestic cat sample (cat IDs: HK29, CB7, OY20, and OI32) (16) and DNA samples from domestic cats in countries, namely, South Korea, Vietnam, Sri Lanka, Tanzania, Japan, and Spain that have been described previously (17).

### Cell cultures

The cells used in this study were cultured in high-glucose Dulbecco’s modified Eagle’s medium (DMEM FUJIFILM; Wako Pure Chemical Corporation, Osaka, Japan) supplemented with 10% fetal calf serum (FCS) and 1X penicillin-streptomycin and incubated in a CO_2_ incubator at 37℃. The following cell types were used: HepG2 (human liver cancer) (18), HEK293T (human embryonic kidney transformed by SV40 large T antigen) (19), AH927 (feline fibroblast cells) (20) CRFK (feline kidney) (21), KwDM (canine mammary gland tumor) (7), Rat2 (RAT-1 [F2408]-cell-derived, thymidine kinase-deficient mutant Rat2 cells) (22), 104C1 (guinea pig, fetal origin) (23), MDTF (*Mus dunni* fibroblast tail) (24), HEK293T cells persistently infected with FeLV-D (293T/FeLV-D cells) (7), 293Lac cells containing a pMXs retroviral vector carrying a LacZ marker (25), and MDTF expressing feline copper transporter 1 (MDTF-feCTR1) cells (26). Briefly, MDTF cells were transfected with expression vectors (pFUΔss-feCTR1 or pFUΔss empty vector as a control) using Lipofectamine®3000 (Thermo Fisher Scientific, Waltham, MA, USA). Single cells were cultured in medium containing 200 μg/mL zeocin (InvivoGen, San Diego, CA, USA) in 96-well plates and single cells were isolated. These cells were designated MDTF-feCTR1 and MDTF-empty for further analysis.

### Cloning of the ERV-DC8 provirus

The DNA genome of domestic cats (cat IDs: HK29, CB7, OY20, and OI32) were used to detect the ERV-DC8 provirus using locus-specific PCR. PCR was performed using KOD One PCR Master Mix (Toyobo, Osaka, Japan), according to the manufacturer’s instructions. Fe-57S (5’-TTAGAGGAATAAGCTCGGGGTAACT-3’) was used as the forward primer, and Fe-54R (5’-GGTGCTCATTGTTAGGAGAGAAAAA-3’) was used as the reverse primer. The cycling conditions were as follows: 35 cycles of denaturation at 98℃ for 10 s, annealing at 60℃ for 5 s, and extension at 68℃ for 100 s. The PCR products were electrophoresed on a 1% agarose gel, stained with ethidium bromide solution, and visualized using a UV transilluminator. The amplified PCR fragment was cloned using pCR® 4 Blunt-TOPO (Invitrogen, Waltham, MA, USA). ERV-DC8 OI32 (cat ID: OI32), which carries a heterologous ERV-DC8, was the ERV-DC8 clone used. The resulting plasmids were confirmed using sequencing (Fasmac Corporation, Atsugi, Japan).

### Construction of the ERV-DC8TA mutant

Sequence analysis and alignment with ERV-DC were performed as previously described (7). A mutant ERV-DC was constructed with thymine (T) replaced with an adenine (A) at nucleotide positions 280 in the 5’LTR and 8594 in the 3’LTR of ERV-DC8 (OI32). The ERV-DC8TA mutant was constructed by site-directed mutagenesis using KOD One PCR Master Mix (Toyobo, Osaka, Japan) with the primers pair DC8-mu6S (5′-CTCCAAGTTGCATCAGCCGAGAGAAACTCC-3′) and DC8-mu6R (5′-GGAGTTTCTCTCGGCTGATGCAACTTGGAG-3′).

### Virus preparation

The plasmids were transfected into 293Lac cells using TransIT®-293 Transfection Reagent (Takara, Shiga, Japan). After 3 days, the supernatant was collected and filtered through a 0.22 µm filter (Merck, Darmstadt, Germany) to prepare a viral stock, and stored at -80℃ for further experiments.

### Infection assay

Approximately 3 × 10^5^ cells/well were seeded in 24-well plates one day before infection. Each well was inoculated with 250 µL of the virus in the presence of 8 μg/mL polybrene (Nacalai Tesque, Kyoto, Japan) for 2 hours. After adding fresh medium, the cells were cultured for 3 days post-infection. For the LacZ assay, culture supernatants were removed and cells were fixed with 250 μL of 2% glutaraldehyde for 15 minutes at room temperature (20–25°C), followed by the addition of 250 μL of 5-bromo-4-chloro-3-indolyl-β-D-galactopyranoside (X-gal) solution to stain cells. After incubation at 37℃ for 2 hours, the nuclei in LacZ-positive cells were counted under a microscope. Viral titers were expressed as infectious units (IU) per mL. The standard deviations were calculated.

### Viral protein preparation

Plasmids were transfected into 293T cells using TransIT®-293 Transfection Reagent. Transfected ERV-DC10 cells were used as positive controls, whereas non-transfected 293T cells were used as negative controls. The cell lysates were prepared by collecting cell pellets 48 and 72 hours after transfection and adding them to cell lysis buffer (20 mM Tris-HCl at pH 7.5, 150 mM NaCl, 10% glycerol, 1% NP-40, 2 mM EDTA, 1 mM Na3VO4, 1 μg/mL aprotinin, and 1 μg/mL leupeptin) on ice for 20 minutes and centrifuging at 15,400 × *g* for 20 minutes at 4 ℃. The protein concentrations were calculated using a protein assay kit (Bio-Rad, Hercules, CA, USA).

For viral purification, cell supernatants were collected 48 and 72 hours after transfection. The supernatant (4 mL) was ultracentrifuged at 29000 × *g* for 90 minutes at 4℃, and the virus pellet was collected in phosphate-buffered saline (PBS) and stored at -80℃ until use.

### Western blotting

To obtain a sample from protein purification, sample buffer (6×) (Nacalai Tesque, Kyoto, Japan) containing 2-mercaptoethanol (FUJIFILM Wako Pure Chemical Corporation) was added and heated at 95-100°C for 5 minutes. Subsequently, SDS-PAGE was performed using a 7.5% gel (Oriental Instruments Co., Ltd.; Sagamihara, Japan) or 4–20% gel (Invitrogen, Carlsbad, CA, USA) at 100 V for 2 hours. The separated protein bands were detected using a primary goat anti-FeLV gp70 polyclonal antibody (National Cancer Institute [NCI], Frederick, MD, USA; dilution 1:15000) as the primary antibody, which cross-reacts with ERV-DC Env and mouse monoclonal anti-human β-actin (Santa Cruz Biotechnology, Dallas. TX). The secondary antibodies used were horseradish peroxidase (HRP)-conjugated donkey anti-goat IgG (Santa Cruz Biotechnology; 1:10000) or HRP-conjugated anti-mouse IgG (Cell Signaling Technology, Danvers, MA, USA). LumiGLO @ Reagent (20×) and 20× peroxide (Cell Signaling Technology, Danvers, MA, USA) were used as substrates for signaling using a Lumino Image Analyzer LAS2000 (Fujifilm, Tokyo, Japan).

### Transmission electron microscopy (TEM)

HEK293T cells transfected with the ERV-DC8TA plasmid were fixed at room temperature for 10 minutes by adding 2% glutaraldehyde in PBS, and then further fixed at 4 ℃ for 10 minutes. Viral particles were analyzed using a transmission electron microscope at the Hanaichi Electron Microscope Technology Research Institute Co., Ltd. (Aichi, Japan).

### ERV-DC8 insertional polymorphism

Chromosomal DNA from domestic cats from Vietnam, Sri Lanka, Tanzania, and South Korea (17) was used for genotyping PCR using the KOD One PCR Master (Toyobo, Osaka, Japan) and the following ERV-DC8 genotyping primers: Fe-57S (5’-TTAGG AATAAGCTCGGGGTAACT-3’), Fe-36S (5’-AACCGCTTGGTACA RTTCATAAGAG-3’), and Fe-54R (5’-GTAGGCTCATTGTTAGGAGAGAAAAA-3’). PCR was carried out using the following cycling conditions: 30 cycles of denaturation at 98℃ for 10 seconds, annealing at 60℃ for 5 seconds, and extension at 68℃ for 6 seconds. The resulting PCR products were analyzed by electrophoresis on a 1% agarose gel.

### Statistical analysis

Infection assay results and geographical correlations were analyzed using one-way analysis of variance, Student’s t-test, or Fisher’s exact test. Statistical significance was set at *p*<0.05.

### Ethical approval

Animal studies were conducted in accordance with the guidelines for the care and use of laboratory animals of the Ministry of Education, Culture, Sports, Science, and Technology, Japan. All the experiments were approved by the Genetic Modification Safety Committee of Yamaguchi University, Japan.

### Accession numbers

The nucleotide sequences reported herein were deposited in the DDBJ, EMBLE, and GenBank databases under the accession number LC597234.

## Results

### ERV-DC8 is infectious and replication-competent

PCR was performed using chromosomal DNA from four domestic cats (cat ID: HK29, CB7, OY20, and OI32) (16), all of which carry ERV-DC8 at the heterologous locus. An approximately 9-kb band corresponding to the ERV-DC8 provirus was detected in all cats, and the ERV-DC8 proviruses from the four cats were molecularly cloned. A preliminary study was conducted using 293Lac and HEK293T cells to determine viral infectivity and replication. All clones were found to be infective; therefore, the ERV-DC8 (OI32) clone was used in this study.

The infectivity of the ERV-DC8 (OI32) virus was evaluated in various cell lines by producing it in 293Lac cells. The ERV-DC8 provirus was transfected into 293Lac cells and the supernatant was used as the viral source. HepG2 (human), KwDM (dog), Rat2 (rat), and 104C1 (guinea pig) cells exhibited viral infectivity ranging from approximately 10^1^–10^3^ infection units (IU)/mL. Infectivity was the highest in HepG2 cells. In contrast, no infection was observed in MDTF (mouse) cells (Figure 1A). Western blotting was performed to detect viral proteins in cell lysates from 293T cells transfected with ERV-DC8. Western blotting yielded a 75-kD protein band corresponding to the Env of ERV-DC8 (Figure 2A). Similarly, purified virions from the cell supernatants also revealed a 75-kD protein band corresponding to Env of ERV-DC8 (Figure 2B). The polyclonal antibody used in this study was sensitive to ERV-DC10 detection. Overall, these findings indicated that ERV-DC8 is an infectious and replication-competent provirus.

**Figure 1.**
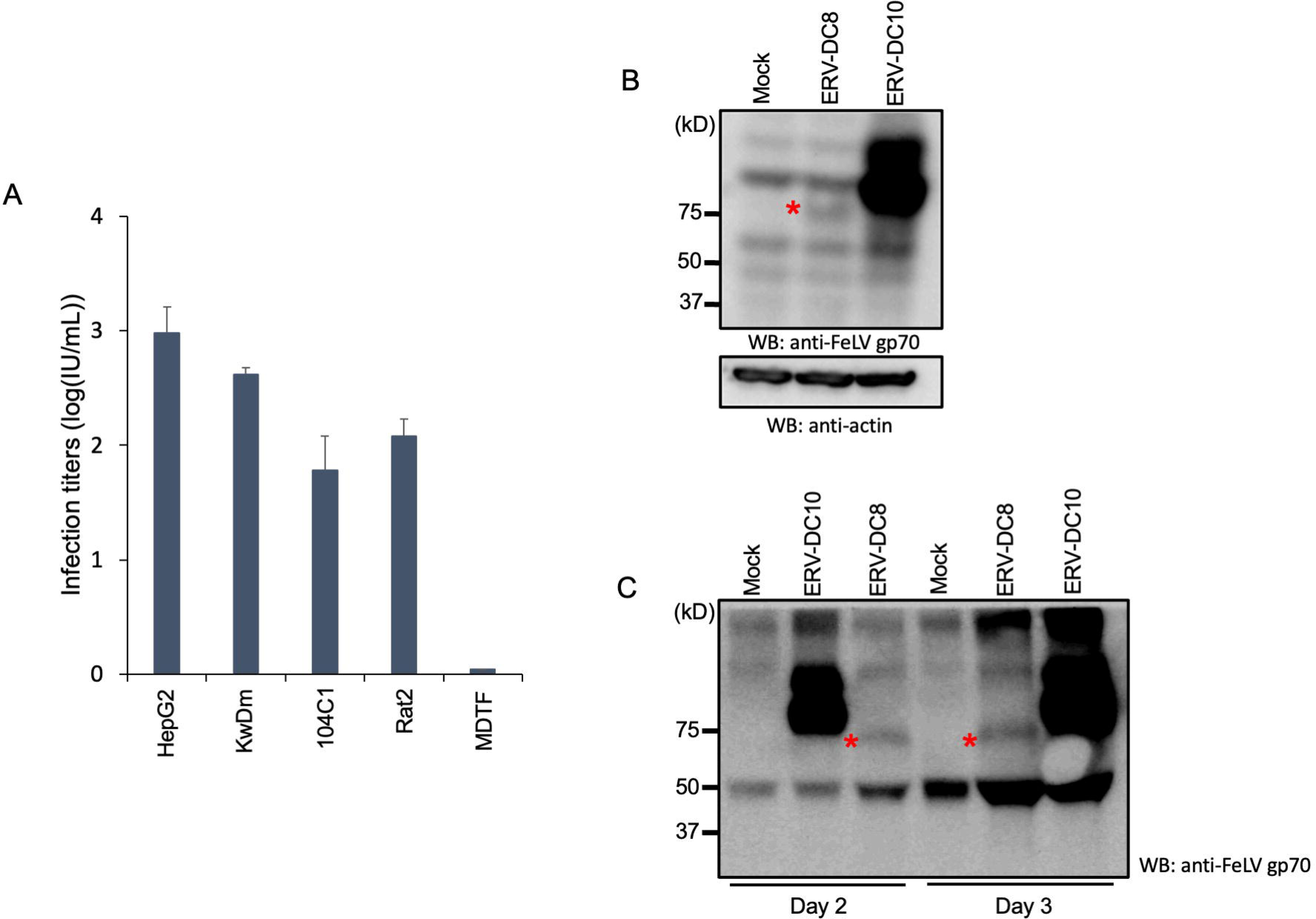
Infection assay of ERV-DC8. (A) HepG2 (human), KwDM (dog), Rat2 (rat), 104C1 (guinea pig), and MDTF (mouse) as target cells were inoculated with ERV-DC8 replication-competent virus. The infectious units were determined by counting the number of log10-galactosidase (LacZ)-positive cells per milliliter of virus. Mean virus infection titers with standard deviations were determined from three independent experiments. (B) Western blot analysis of ERV-DC Envs proteins in 293Lac cells using anti-FeLV gp70. Actin was used as a control. (C) ERV-DC8 Env protein was detected in purified virions from the supernatant of HEK293T cells transfected with ERV-DC8 at 2 and 3 days post-transfection. ERV-DC10 was used as the positive control. Mock cells were used as the negative control. Anti-FeLV gp70 was used for western blotting, whereas anti-human β-actin was used as the control. Red asterisks indicate Env proteins.

**Figure 2.**
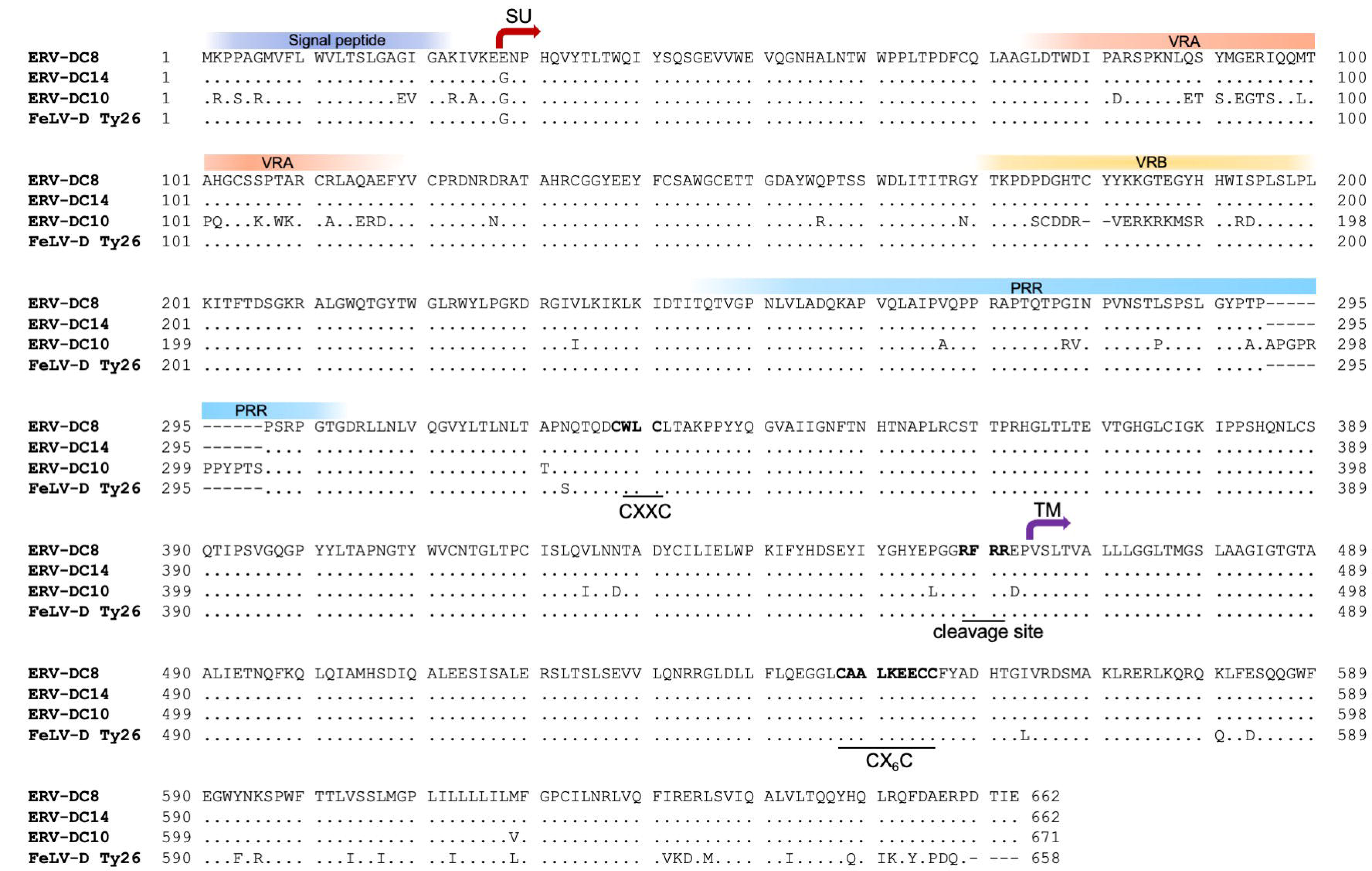
Amino acid sequence alignment of ERV-DC8 Env. Amino acid sequence alignment of the Env proteins of ERV-DC8 (in this study; LC597234), ERV-DC14 (AB674445), ERV-DC10 (AB674444), and FeLV-D TY26 (AB673428). SP, signal peptide; SU, surface unit; VRA, variable region A; VRB, variable region B; PRR, proline-rich region; TM, transmembrane subunit; R-X-R/K-R is the cleavage motif; CXXC and CX6CC are sites of covalent interaction.

### Sequence analysis of ERV-DC8 (OI32)

The ERV-DC8 (OI32) nucleotide sequence was 8864 bp in length, consisting of a 5244 bp *gag-pol* and a 1990 bp *env* gene, and an intact ORF (Figure 2), similar to the previously isolated ERV-DC8 IS10 clone, which was not replication-competent. A comparison of the ERV-DC8 OI32 and IS10 sequences revealed a one-base difference at position 3162 in the *pol* gene. ERV-DC8 OI32 had a guanine (G), whereas ERV-DC8 IS10 had a T at this position. ERV-DC8 from cats ID HK29, CB7, and OY20 also contained guanine (G). The ERV-DC8 Env protein is composed of 662 amino acids and contains a putative signal peptide (SP), SU subunit, furin cleavage site, and TM subunit. It is structurally identical to ERV-DC14 Env, with only one amino acid difference in the SU subunit at position 28. Additionally, ERV-DC8 shared a 96.5% amino acid sequence with FeLV-D/Ty26 Env (Figure 2).

### Single nucleotide polymorphism (SNP) in the LTRs of ERV-DC8

The nucleotides at positions 280 in the 5’LTR and 8594 in the 3’LTR of ERV-DC8 (OI32) were both T, indicating that the SNP affected the LTR promoter activity in ERV-DC (25). Therefore, an ERV-DC8TA mutant, in which T was replaced with A, was constructed and tested for infectivity and viral expression (Figure 3A). The viral titer of the ERV-DC8TA mutant was approximately 4-fold higher than that of ERV-DC8 in HEK-293T cells (Figure 3B). Furthermore, ERV-DC8 Env protein expression was higher in ERV-DC8TA than in wild-type ERV-DC8 (Figure 3C). ERV-DC14 and ERV-DC14TA were used as controls as previously described (25). Furthermore, ERV-DC8TA inoculation into feline cell lines (AH927 and CRFK) let to infectivity in both cell lines (Figure 3D). Our results suggest that the LTR promoter activity in ERV-DC8 is affected by a single nucleotide change from T to A.

**Figure 3.**
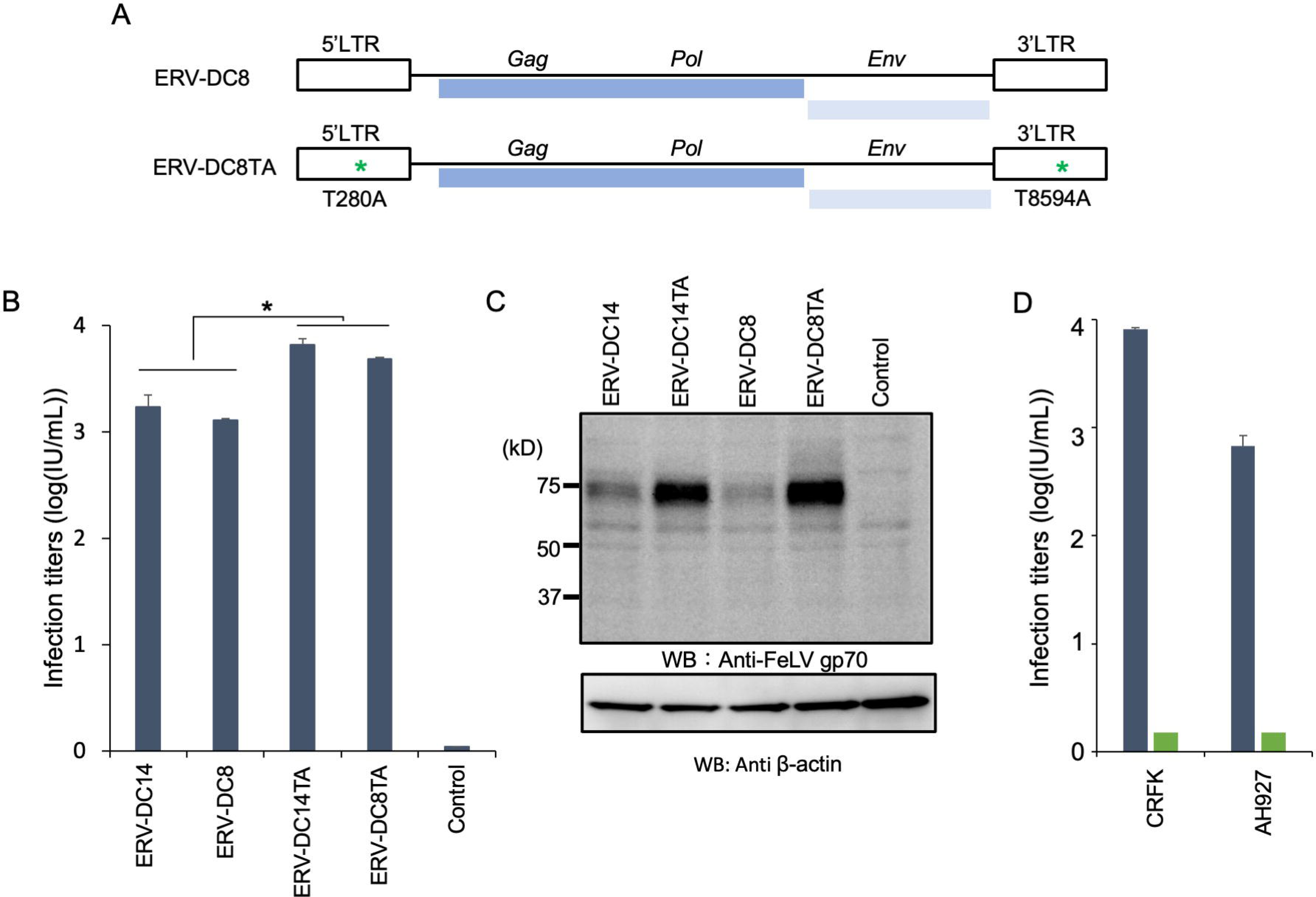
Analysis of the ERV-DC8TA mutant. (A) Schematic representation of the ERV-DC8 and ERV-DC8TA mutants. Green asterisks indicate nucleotide position. Thymine (T) was replaced with adenine (A) at nucleotide positions 280 in the 5’LTR and 8594 in the 3’LTR of ERV-DC8 (OI32). (B) Infection assay of the ERV-DC8TA mutant. HEK293T (human) cells were infected with the indicated viruses. (C) Western blot analysis of the ERV-DC8TA mutant. The cell lysates from HEK293T cells transfected with the indicated ERV-DCs were used to detect ERV-DC8 Env protein 3 days after transfection. Anti-FeLV gp70 was used for western blotting, whereas anti-human β-actin was used as the control. (D) Infection assay of the ERV-DC8TA mutant in feline cell lines, AH927 and CRFK cells. The infectious units were determined by counting the number of log10-galactosidase (LacZ)-positive cells per milliliter of virus. Mean virus infection titers with standard deviations were determined from three independent experiments. The data were statistically analyzed by one-way analysis of variance (\**p*<0.0001).

### Transmission electron microscopy (TEM) analysis

TEM analysis of HEK293T cells infected with ERV-DC8TA revealed 90–100-nm viral particles (Figure 4). This result suggests the viral budding morphology, characterizing the virus as a type C retrovirus or *Gammaretrovirus*.

**Figure 4.**
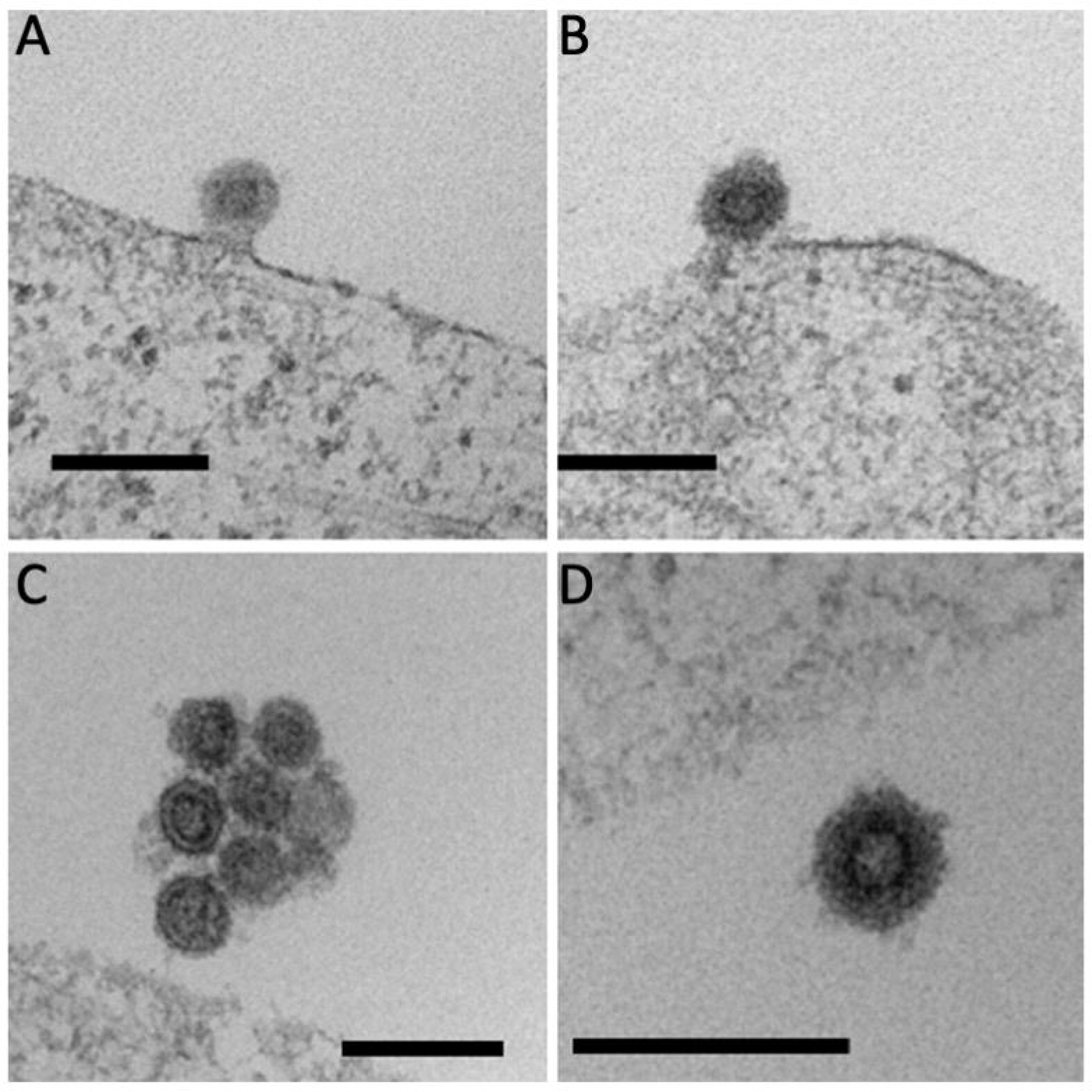
Transmission electron microscopy (TEM). HEK293T cells transfected with ERV-DC8TA mutant were cultured for 1 week, fixed, and analyzed by TEM. (A, B) The cells demonstrated viral budding, and (C, D) the viral particles. The scale bar indicates a length of 200 nm.

### Receptor for ERV-DC8

Previous reports have demonstrated that the ERV-DC8 Env-pseudotyped virus is in the same interference group as FeLV-D (27). In this study, we performed an interference assay using the ERV-DC8 OI32 replication-competent virus. The viral titer of ERV-DC8 was found to be 3.3 × 10^2^ IU/mL when HEK293T/mock cells were infected, compared to 1.3×10^0^ IU/mL when HEK293T/FeLV-D cells were infected. The ERV-DC8 virus significantly interfered with FeLV-D infection (*p*<0.05). FeLV-D and ERV-DC genotype I have previously been reported to use feCTR1 as a receptor (26, 28). To confirm the receptor for ERV-DC8, the virus was inoculated into MDTF-feCTR1 cells. MDTF-feCTR1 permitted ERV-DC8 infection (Figure 5B). Overall, these findings indicate that ERV-DC8 uses the same receptor as FeLV-D, that is, the feCTR1.

**Figure 5.**
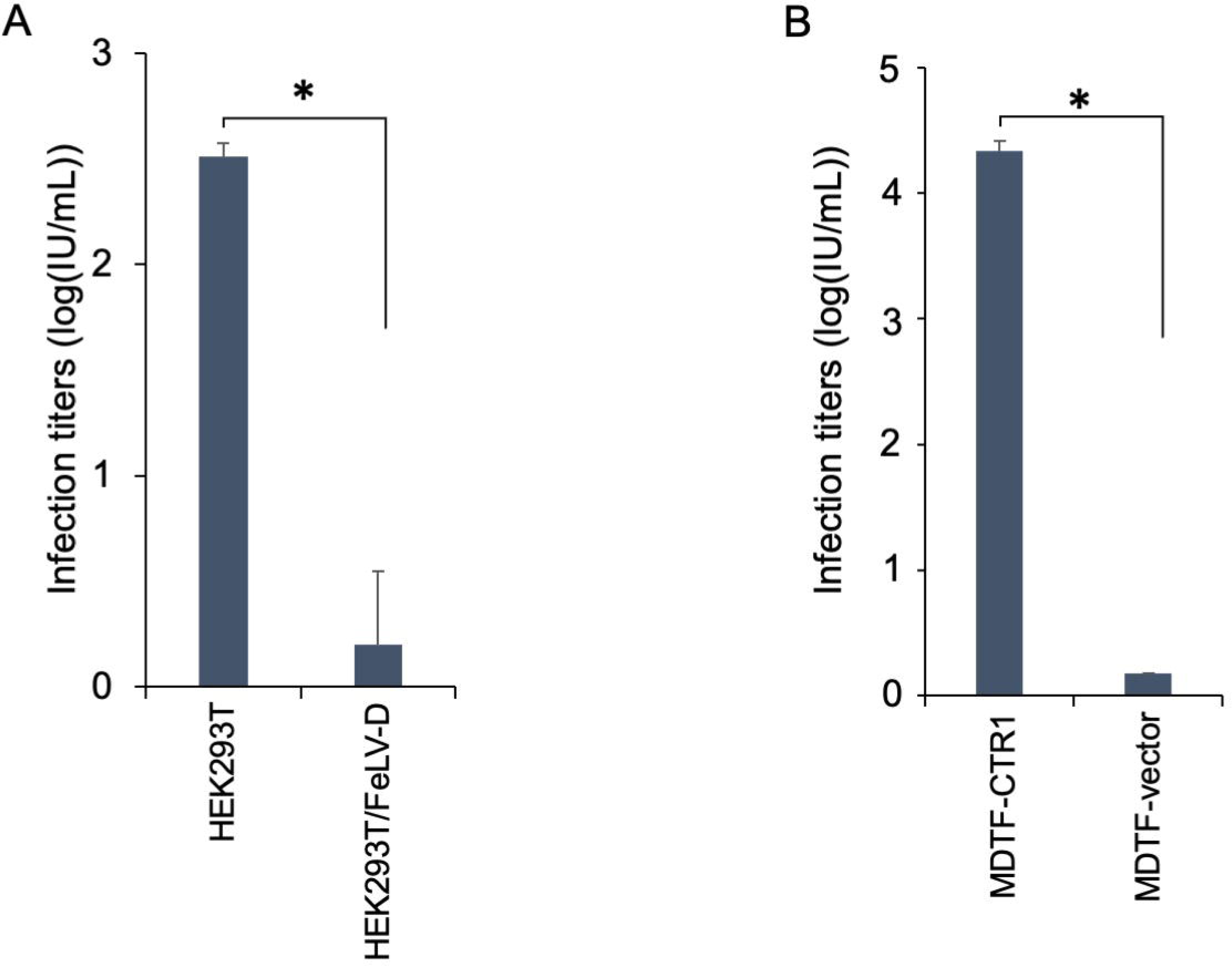
Interference assay with FeLV-D and receptor usage of ERV-DC8. (A) HEK293T cells persistently infected with FeLV-D were inoculated with ERV-DC8 containing a retrovirus vector carrying a LacZ marker. (B) MDTF-CTR1 cells were inoculated with ERV-DC8TA mutant carrying a LacZ marker. MDTF-vector indicates the control. The infectious units were determined by counting the number of log10-galactosidase (LacZ)-positive cells per milliliter of virus. Mean virus infection titers with standard deviations were determined from three independent experiments. The data were statistically analyzed by student-t test (\**p*<0.005).

### Insertional polymorphism of ERV-DC8

Previously, we investigated the prevalence of ERV-DC8 in domestic cats in Japan (*n* = 244) and Spain (*n* = 35) (29). Our findings revealed a high frequency of ERV-DC8 in domestic cats from Japan (77.5%) and Spain (65.7%) (7, 29). Here, we further investigated the frequency of ERV-DC8 in domestic cats using PCR primers based on unique genomic DNA flanking sequences or with secondary primers based on the proviral sequence (Figure 6A). PCR revealed that the provirus was not present on either chromosome (indicated by the −/− symbol), was present as homozygous (indicated by the +/+ symbol), or was present as heterozygous (indicated by the +/− symbol) (Figure 6B). PCR was performed to detect ERV-DC8 in domestic cats from Tanzania (*n* = 49), Vietnam (*n* = 19), South Korea (*n* = 19), and Sri Lanka (*n* = 35). The frequency of ERV-DC8 ranged from 52.6% to 85.8% in domestic cats from these four countries (Figure 6C). The frequency of ERV-DC8 was the highest among domestic cats in Tanzania (85.8%) and the lowest among domestic cats in Vietnam (52.6%) and South Korea (52.6%) (Figure 6C). The frequency of ERV-DC8 in Asia was the highest among domestic cats in Japan (29), with similar proportions observed in Vietnam and South Korea. The prevalence of ERV-DC8 varied significantly among the domestic cats in the countries studied. There were significant differences between Tanzania and Sri Lanka (*p*=0.0375), Tanzania and South Korea (*p*=0.0087), Tanzania and Vietnam (*p*=0.0087), Vietnam and Japan (*p*=0.024), and South Korea and Japan (*p*=0.024). Collectively, these results suggest that ERV-DC8 is highly integrated into domestic cats in these countries.

**Figure 6.**
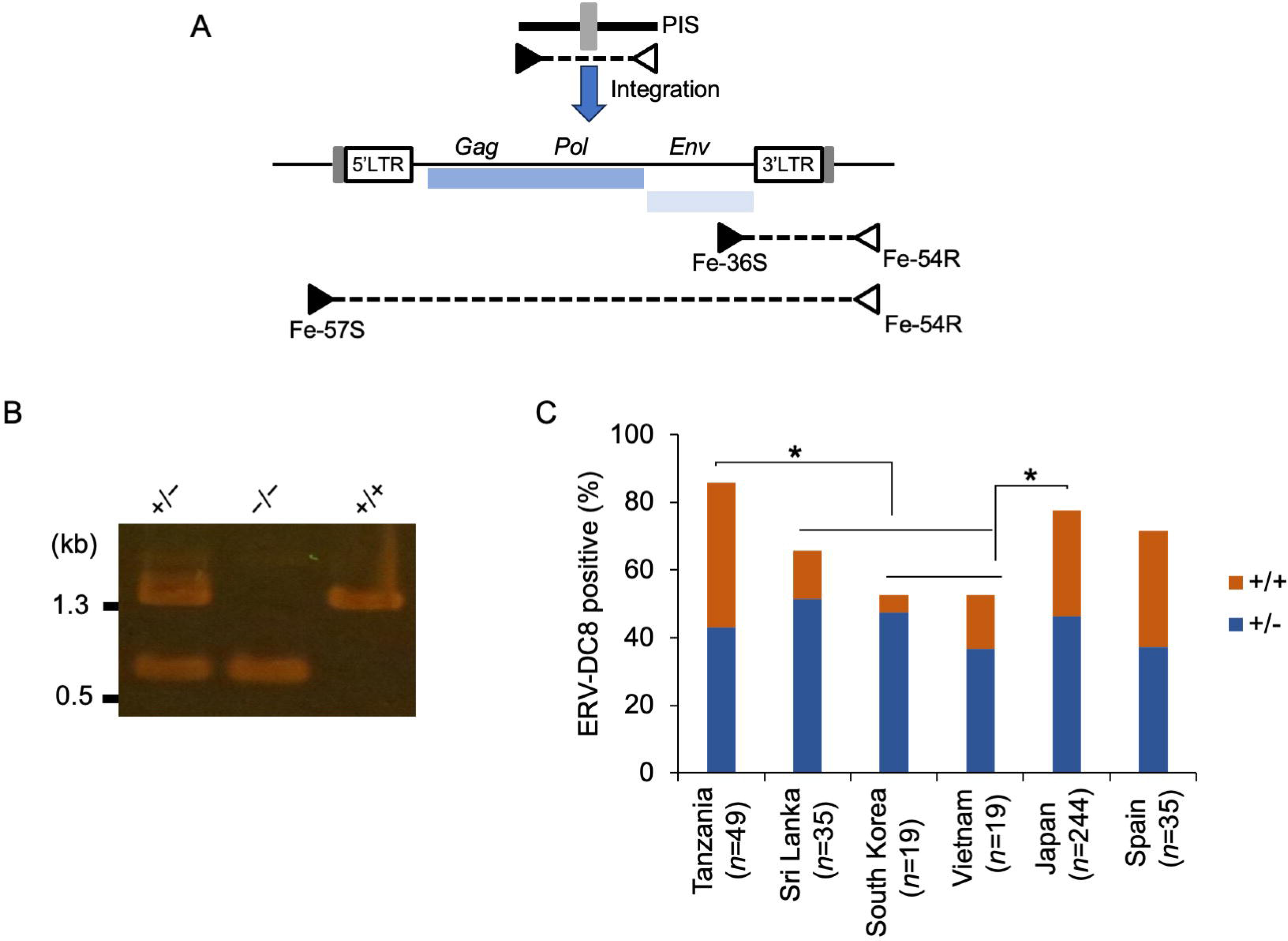
Insertional polymorphism of ERV-DC8. (A) PCR for pre-integration site (PIS) detection and provirus using the indicated PCR primers. Primer pairs Fe-57S and Fe-54R detected a PIS and/or full-length provirus. (B) ERV-DC8 detection and genotyping. Band sizes of 0.5 kbp and 1.3 kbp represent the pre-integration site and proviral insertional polymorphic sites, respectively. −/−, no copy of the provirus present on either chromosome; +/+, proviral copy present on both chromosomes (homozygous); +/−, provirus copy present on one of two chromosomes (heterozygous). (C) Prevalence of ERV-DC8 in domestic cats from different countries. The frequencies of ERV-DC8 in domestic cats from different countries. +/+, copy present on both chromosomes (homozygous, orange); +/−, copy present on one of two chromosomes (heterozygous, blue). The data were analyzed using Fisher’s exact test (\**p*<0.05). Data from Japan and Spain was obtained from a previous report (29).

## Discussion

In this study, we characterized ERV-DC8 by examining its genetic structure, sequence differences from related ERVs, and infectious abilities. We established ERV-DC8 as infectious and replication-competent. Recently four ERV-DCs have been identified as infectious and replication-competent: ERV-DC8 (in this study; LC597234) and ERV-DC14 (25) are classified as GI, and ERV-DC10 and ERV-DC18 are classified as GIII (7). Only one cat family has been shown to harbor ERV-DC18, which contains a single nucleotide difference compared to ERV-DC10, suggesting that it might be internalized via reinfection with ERV-DC10 (7). ERV-DC8 and ERV-DC14 differ in sequence by 24 nucleotides. Both ERV-DC8 and ERV-DC14 exhibit a single nucleotide difference in the sequences of the 5’ and 3’LTRs. Therefore, sequence analysis indicates that ERV-DC is a young ERV (7, 15). Recent studies have shown that an SNP exists in the Env of ERV-DC14 in European wildcats (*Felis silvestris*), resulting in replication incompetence (29). Our western blot analysis and viral infection assays (Figure 3) showed an increased level of replication of the ERV-DC8TA mutant compared to that of the wild type due to the change in promoter activity (Figure 4B). This phenomenon is similar to that observed in ERV-DC14, and the attenuated promoter activity may be a mechanism of ERV inactivation.

In investigating the ability of the ERV-DC8 provirus to infect various hosts, we found that ERV-DC8 provirus carrying LacZ infected cells derived from humans (HEK293T and HepG2), cats (AH927 and CRFK), dogs (KwDM), rats (Rat2), and guinea pigs (104C1), but not from mice (MDTF). The previously cloned ERV-DC8 (IS10) was replication-incompetent because of a nucleotide change at position 3162 in the reverse transcriptase gene (7), which we compared with our clone ERV-DC8 (OI32) in this study.

Our findings provide valuable insights into the insertional polymorphism of ERV-DC8. ERV-DC8 insertional polymorphism frequency ranged from 53–86% in domestic cats from several countries (Japan (29), Spain, Tanzania, Vietnam, South Korea, and Sri Lanka). Although a geographical relationship could not be determined with certainty in this study, ERV-DC8 is spreading among domestic cats worldwide. However, ERV-DC8 was not integrated into European wildcats (29). We hypothesize that ERV-DC8 may not have integrated into the same ancestral line of cats. The integrated ERV-DC8 ancestor may have been present in the lineage of domestic cats but absent in that of wildcats lineage.

Additionally, a viral interference assay revealed that infectious ERV-DC8 uses the same receptors as FeLV-D, which was confirmed by infection in MDTF-feCTRI cells. FeLV-D is generated using the *env* genes of FeLV and the recombination of ERV-DC GI to acquire a new viral property (7, 15). FeLV-D is associated with lymphoma/leukemia induction (12). We speculate that ERV-DC8 may represent a major contributor to the emergence of FeLV-D because of the high frequency of ERV-DC8 in domestic cats.

Several animals such as cats and pigs have been reported to have infectious ERVs (7, 30), which is in line with our findings of ERV-DC8 in domestic cats. Activation of infectious ERVs is potentially harmful to the body. The presence of infectious ERVs in domestic cats can be dangerous to the host because they contribute to the emergence of several FeLV subtypes (12). Mice have also been reported to have infectious ERVs, which can lead to tumors in patients with TLR7 deficiency or immunodeficiency, owing to ERV activation (31). APOBEC, an antiretroviral molecule, suppresses the emergence of pathogenic ERVs (32). Thus, host immunity and defense mechanisms regulate ERV activation. Furthermore, even in the presence of such ERV, epigenetic control prevents the virus from being expressed by losing its transcriptional activity (4). In contrast, some ERVs have been reported to benefit the host; for example, Syncytin-1 aids placentation (33), Refrex-1 promotes cellular copper homeostasis (34), and resistance conference against retrovirus infection in several species, including cats and primates (26). The existence of ERVs in hosts can be attributed to several factors—harmful and beneficial.

In conclusion, our findings establish the existence of infectious ERV-DC8 in domestic cats. ERV-DC8 may play a major role in the emergence of FeLV-D, which is pathogenic to the host (cats). ERV-DC8 dynamics must be further studied for feline health management. This study also highlighted the role of ERVs and the evolutionary relationship between viruses and their hosts. Future research should focus on elucidating the mechanisms of ERV activation and its impact on host immunity to develop strategies for mitigating risks while harnessing potential benefits.

## Acknowledgments

We are grateful to Dr. Yoshinao Kubo for providing the MDTF cells and Dr. Toshio Kitamura for providing the GP cells and pMxs retroviral vector. We thank Editage (www.editage.jp) for the English language editing.

## Author contributions statement

Conceptualization: KN; Data Curation: DP and YM; Formal analysis: DP, YM, and KN; Funding acquisition: KN; Investigation: DP, YM, YS, RMCD, IM, MHN, DFL, AM, and KN; Methodology: DP, YM, and KN; Project administration: KN; Supervision: KN; Validation: DP, YM, and KN; Writing-original draft: DP and KN; Writing-review & draft: DP and KN.

